# *Lactobacillus rhamnosus* attenuates bone loss and maintains bone health by skewing Tregs-Th17 cell balance in Ovx mice

**DOI:** 10.1101/2020.08.19.257048

**Authors:** Leena Sapra, Hamid Y. Dar, Amit Pandey, Surbhi Kumari, Zaffar Azam, Asha Bhardwaj, Prashant Shukla, Pradyumna K. Mishra, Bhupendra Verma, Rupesh K. Srivastava

## Abstract

Osteoporosis is a systemic-skeletal disorder characterized by enhanced fragility of bones leading to increased rates of fractures and morbidity in large number of populations. Probiotics are known to be involved in management of various-inflammatory diseases including osteoporosis. But no study till date had delineated the immunomodulatory potential of *Lactobacillus rhamnosus* (LR) in bone-health. In the present study, we examine the effect of probiotic-LR on bone-health in osteoporotic (Ovx) mice model. We observed that administration of LR attenuated bone-loss in Ovx mice. Both the cortical and trabecular bone-content of LR treated group was significantly higher than Ovx-group. Remarkably, the percentage of osteoclastogenic-CD4^+^Rorγt^+^Th17 cells at distinct immunological sites such as BM, spleen, LN and PP were significantly reduced, whereas the percentage of anti-osteoclastogenic-CD4^+^Foxp3^+^Tregs and CD8^+^Foxp3^+^Tregs were significantly enhanced in LR-treated group thereby resulting in inhibition of bone-loss. The immunomodulatory-role of LR was further supported by serum-cytokine data with a significant reduction in proinflammatory-cytokines (IL-6, IL-17 and TNF-α) along with enhancement in anti-inflammatory-cytokines (IL-10, IFN-γ) in LR treated-group. Altogether, the present study for the first time establishes the osteoprotective role of LR on bone-health, thus highlighting the potential of LR in the treatment and management of various bone related diseases including osteoporosis.

## Introduction

Bone is a dynamic organ that maintains its proper architecture and function by undergoing continuous cycles of modelling and remodelling and thus, helps in maintaining normal host physiology. Any dysregulation in the bone-remodelling process results in development of bone related diseases including osteoporosis. Patients diagnosed with osteoporosis exhibit low mineral-density along with deteriorating micro-architecture of bones and are thus at higher risk of fractures (1–4). Among the global population, greater prevalence of osteoporosis has been observed in postmenopausal women due to lower estrogen levels (5). Various FDA approved drugs and monoclonal antibodies are being currently used for treatment of osteoporosis, but unfortunately these compounds along with providing relief to patients also cause various side effects (6). Due to increase in ageing population across the globe, osteoporosis is now becoming a budding medical and socioeconomic issue worldwide and thus there is an exigent need to identify safe and cost effective interventions exhibiting both preventative and therapeutic abilities for management of osteoporosis.

Estrogen hormone is an important regulator of bone density. The progression towards osteoporosis is accelerated at faster rate in postmenopausal women due to the declining levels of osteoprotective hormone “estrogen”. The significance of estrogen hormone in maintaining bone homeostasis is illustrated by the fact that deficiency of hormone leads to the development of postmenopausal osteoporosis in women (7). One of the crucial downstream mediators of estrogen action in maintaining the bone homeostasis is osteoprotegerin (OPG)/ receptor activator of NFkB ligand (RANKL) system (8). RANKL is a crucial cytokine involved in differentiation of osteoclasts precursors and in survival of mature osteoclasts. RANKL is expressed by variety of cells such as osteoblasts, T cells and B cells (9).

For past few years, Tregs and Th17 cells have gained tremendous attention due to their association with various autoimmune and inflammatory diseases (10). CD4^+^Foxp3^+^Tregs cell via secreting anti-inflammatory or immunosuppressive cytokines such as IL-10 and TGF-β suppresses osteoclastogenesis and bone resorption in a cytokine dependent manner. On the contrary, CD4^+^ Rorγt^+^ Th17 cell via secreting IL-6, IL-17 and TNF-α inflammatory cytokines enhance the expression of RANKL on osteoblasts and fibroblasts and thus promotes osteoclasts mediated bone resorption (11). Furthermore, several studies demonstrated that decline in estrogen levels not only effects the bone forming (osteoblastogenesis) and bone resorbing (osteoclastogenesis) process but is also known to affect functionality of immune cells especially T cells (12, 9, 13). Also, the homeostatic balance of Treg-Th17 cell axis is an important determinant of enhanced bone-loss in osteoporosis (3, 14). Estrogen exhibits the potential to stimulate the differentiation and survival of Tregs which have already been shown to suppress bone resorption (15). Tyagi et al. demonstrated that estrogen via suppressing secretion of inflammatory cytokines from Th17 cells suppresses bone resorption (16). Thus, bone-loss associated with estrogen deficiency may occur due to the impairment in complex network of hormones and cytokines that interrupts the bone remodelling process. In lieu of the growing involvement of immune system in osteoporosis, our group had recently coined the term “Immunoporosis” (i.e. Immunology of Osteoporosis) to highlight the specific role of immune system in pathology of osteoporosis (4).

In recent years, “gut-bone” axis has gained tremendous attention of researchers. It has been observed that any dysbiosis in the intestinal microbiota leads to pathogenesis of various diseases such as IBD, obesity, diabetes, RA etc. and all these disease conditions can further lead to bone-loss and development of secondary osteoporosis (17–21). The appreciation that certainly bacterial species can impart numerous benefits to human health dates to ancient times. This has led to tremendous expansion in the field of “Probiotics” in the last 5 years. WHO defines Probiotics as viable microorganisms acting as nutritional supplement that confers various health benefits when administered in adequate amounts (22, 23). The first idea related to beneficial effects of probiotics was suggested in early 20^th^ century by Eli Metchnikoff. Also, several animal and small-scale human studies reported that intake of probiotics showed positive results in Ovx mice and in osteoporotic patients (24, 3, 14, 25).

One of the widely studied probiotic strain of *Lactobacillus* species for various human applications is *Lactobacillus rhamnosus* (LR). It is a gram positive, anaerobic bacterium that exhibits the capacity to transport and metabolize carbohydrates and thereby helps in maintaining the epithelial-layer gut integrity (26). Recently, a study demonstrated that LR-administration alleviates gut inflammation and improved barrier function of intestine (27). LR-administration is also found to enhance bone mass in eugonadic mice (13). Earlier, we too have reported the immunomodulatory properties of LA and BC probiotics in Ovx mice (3, 14). But the immunoregulatory role of LR in regulating bone-health is still required. Thus, in the present study we aim to investigate the immunoregulatory role of LR on bone-health in Ovx mice.

Herein, we report that administration of LR suppresses bone resorption and maintains bone mass in Ovx mice via augmenting the levels of anti-inflammatory cytokines (IL-4, IL-10 and IFNγ) and suppressing the levels of inflammatory cytokines (IL-6, IL-17, RANKL and TNF-α) by maintaining the balance between Tregs-Th17 cells equilibrium in Bone marrow (BM), Spleen and Lymph Nodes (LN). Interestingly, LR also exhibits the ability in modulating the Tregs-Th17 cell balance in the anatomically important intestinal lymphoid tissues Peyer’s patch (PP). Altogether, the present study highlights the specific effect of LR-administration on bone mass by modulating the physiologically important Tregs-Th17 cell balance, thereby opening novel avenues in the management and treatment of various inflammatory bone diseases including osteoporosis.

## Results

### *Lactobacillus rhamnosus* attenuates bone-loss in ovariectomized mice

Recent reports have highlighted the significance of probiotics supplementation in regulating bone-health (3, 14, 13, 25). In present study, we inspected the role of probiotic LR in regulating bone-health via its immunomodulatory properties, since no specific study has been done till date to delink the properties of LR in the regulation of bone-health in Ovx mice. During LR treatment, we firstly examined the body weight of all groups at regular intervals (Day 1^st^, 21^st^ and 45^th^) and we did not observe any significant change in the body weights of mice (Fig. 1b). To investigate whether LR-administration specifically inhibits bone-loss in Ovx mice, we next studied the effect of LR-administration on bone pathology and bone remodelling processes. Analysis of femurs (cortical) bones collected at the end of experiment by SEM revealed that mice of Ovx group had enhanced number of resorption pits or lacunae representing higher osteoclastogenesis as compared to Ovx+LR administrated group where significantly reduced resorption pits/lacunae were found, a clear sign of lower osteoclastogenesis (Fig. 2a). Since SEM images illustrate 2D-information of bone samples and we were also interested to study the 3D-topology of bone-structures, so we next performed AFM analysis of femur cortical bones. Since, AFM finds an impeccable place in structural biology for visualization of biological structures with unique ability to measure molecular architecture with higher sensitivity. We therefore employed AFM for derivation of ultra-structural information of bone surfaces where we observed a significant suppression in bone resorption in LR administered group as compared placebo treated Ovx group (Fig. 2c). This conclusive data from AFM further supplemented and validated our earlier SEM data and thus supports our hypothesis that LR inhibits bone-loss in Ovx mice.

**Fig 1:**
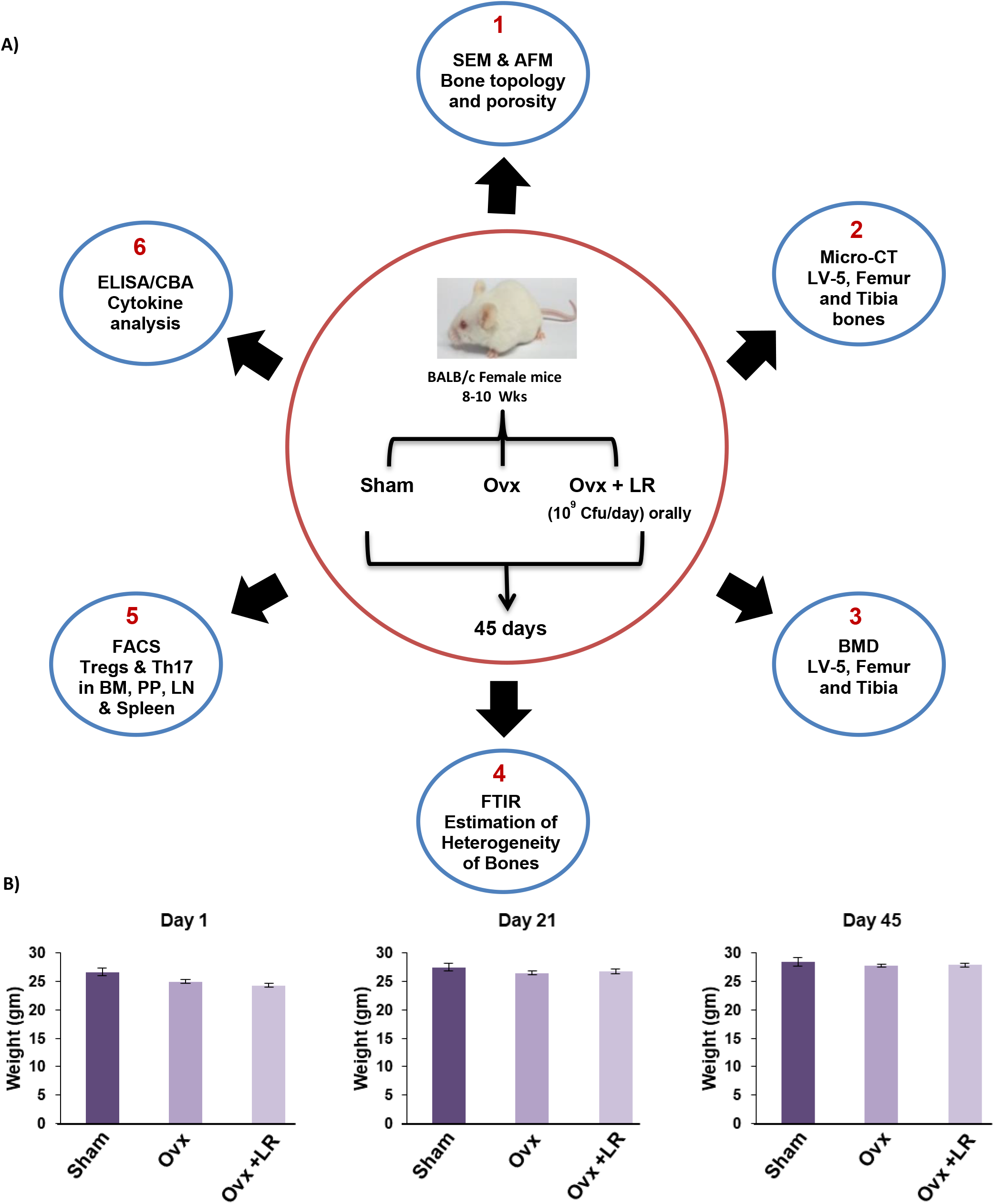
Experimental work plan & Body weight. A) Mice were divided into 3 groups viz. Sham group, Ovx group received placebo control and Ovx + LR group received LR at 109 CFU/day orally reconstituted in drinking water. At the end of 45 days, mice were sacrificed. B) Effect of Lactobacillus rhamnosus on Body Weight; Values are reported as mean ± SEM (n=6). (Mouse Image courtesy: Hamid Y. Dar).

**Fig 2:**
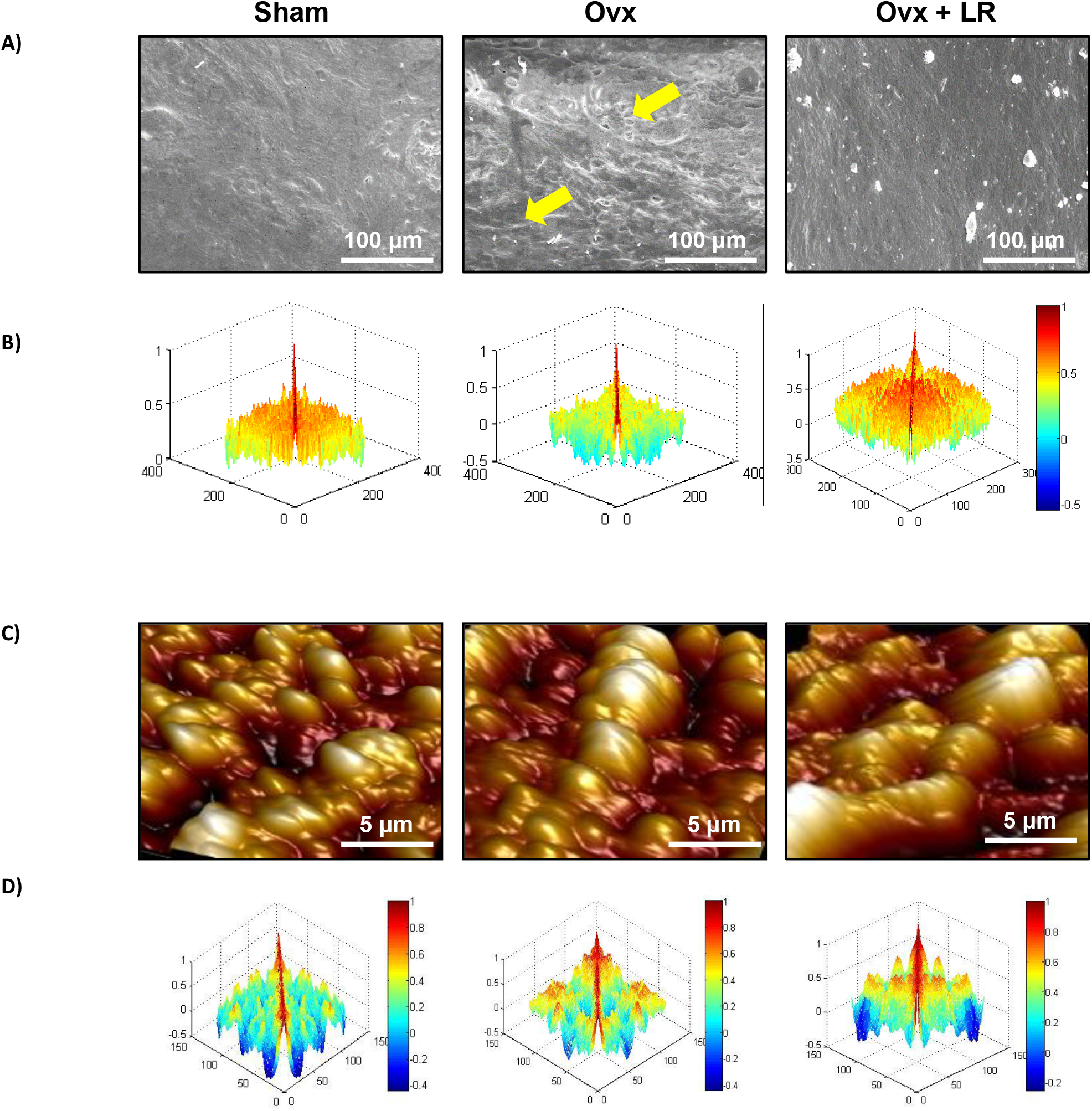
LR administration attenuates bone loss. Mice were sacrificed and cortical bones of all groups were collected for SEM and AFM analysis. A) 2D SEM images. B) 2D MATLAB analysis of SEM images C) 3D AFM images D) 2D MATLAB analysis of AFM image. The above images are indicative of one independent experiment and comparable results were obtained in two different independent experiments with n=6 mice/group/experiment.

To further analyse SEM and AFM images quantitatively in a more statistical way, a well-known computing technology called MATLAB-analysis (matrix-laboratory) was employed to derive the correlation between bone-mass and bone-loss. In MATLAB usually colour representation is used to derive an analogy for evaluating different variables which gives idea about distortion of identical objects with varying correlation values. MATLAB analysis of 2D-SEM images signifies the degree of homogeneity where red colour symbolizes higher correlation (high bone mass) whereas blue colour symbolizes lesser correlation (more-bone-loss). Thus, from the MATLAB-data of SEM (Fig. 2b), we can summarize that Ovx+LR group showed greater correlation and more bone mass. Similarly, MATLAB-analysis of AFM-3D images (Fig. 2d) represented height of the bone where red colour represents enhanced bone architecture (reduced-osteoclastogenesis) and blue colour represents reduced bone-architecture (enhanced-osteoclastogenesis). Thus, it can be inferred from (Fig. 2d) that Ovx+LR group showed more correlation value with enhanced bone architecture or suppressed osteoclastogenesis as compared to Ovx group. Together, findings from both SEM and AFM data analysis specify that LR-administration significantly reduced bone resorption in Ovx mice.

### LR supplementation preserves bone-micro-architecture in Ovx mice

Next, we were interested in studying the effect of LR-administration on the bone micro-architecture of Lumbar-vertebrae-5 (LV-5). We thus performed high resolution μ-CT imaging (a gold standard for determining bone-health) to assess the morphology of trabecular region of LV-5 and also analysed the morphometric parameters that depends on 3D algorithms about the underlying structures but not on the assumptions (28). In bone biology LV-5 region is being considered as one of the most peculiar regions to diagnose early bone-loss or osteoporosis (3, 14, 29). μ-CT 3D images showed that the micro-structure of LV-5 trabecular region of Ovx group was spotted by reduced interconnections within trabecular region along with significantly enhanced space as compared to sham group (Fig. 3). Strikingly, u-CT data clearly demonstrated significant improvement in LV-5 micro-architecture of Ovx+LR group in comparison to Ovx group (Fig. 3). From the μ-CT data it was further found that LR-administration in Ovx mice significantly increased bone volume/tissue volume (BV/TV) (p < 0.05), trabecular thickness (Tb.Th) (p < 0.01) and connective density (Conn.Den) (p < 0.05) (Fig. 4), whereas trabecular separation (Tb.Sp.) (p < 0.05) was found to be significantly reduced as compared to Ovx group. These findings demonstrate and prove that LR-administration in Ovx mice tend to improve the trabecular bone microarchitecture of LV-5 bones.

**Fig 3:**
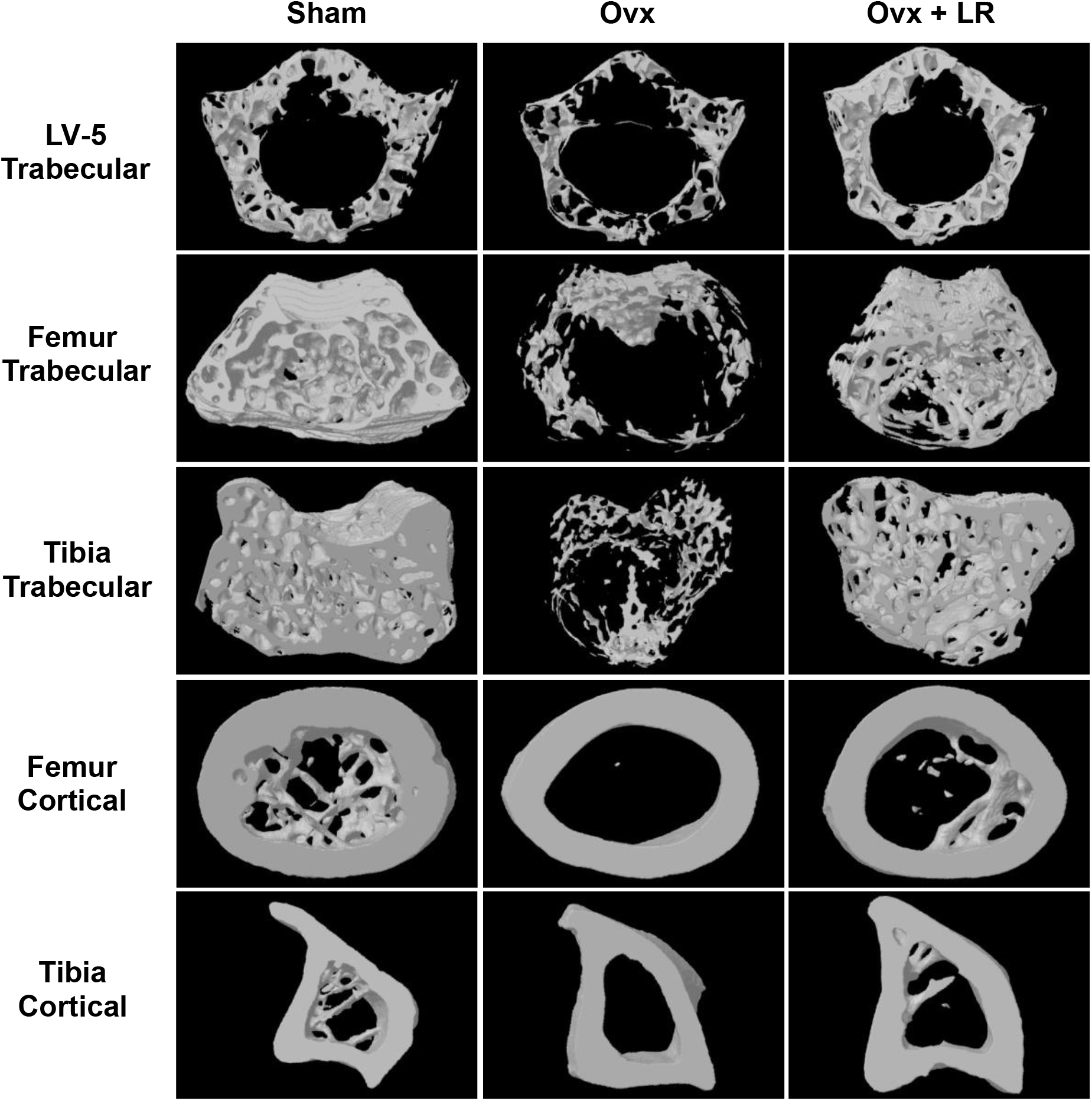
LR administration enhances trabecular and cortical bone microarchitecture. 3-D uCT reconstruction of LV-5 Trabecular, Femur Trabecular, Tibia Trabecular, Femur cortical and Tibia cortical of all groups. Given images are indicative of one independent experiment and comparable results were obtained in two different independent experiments with n=6 mice/group/experiment.

**Fig 4:**
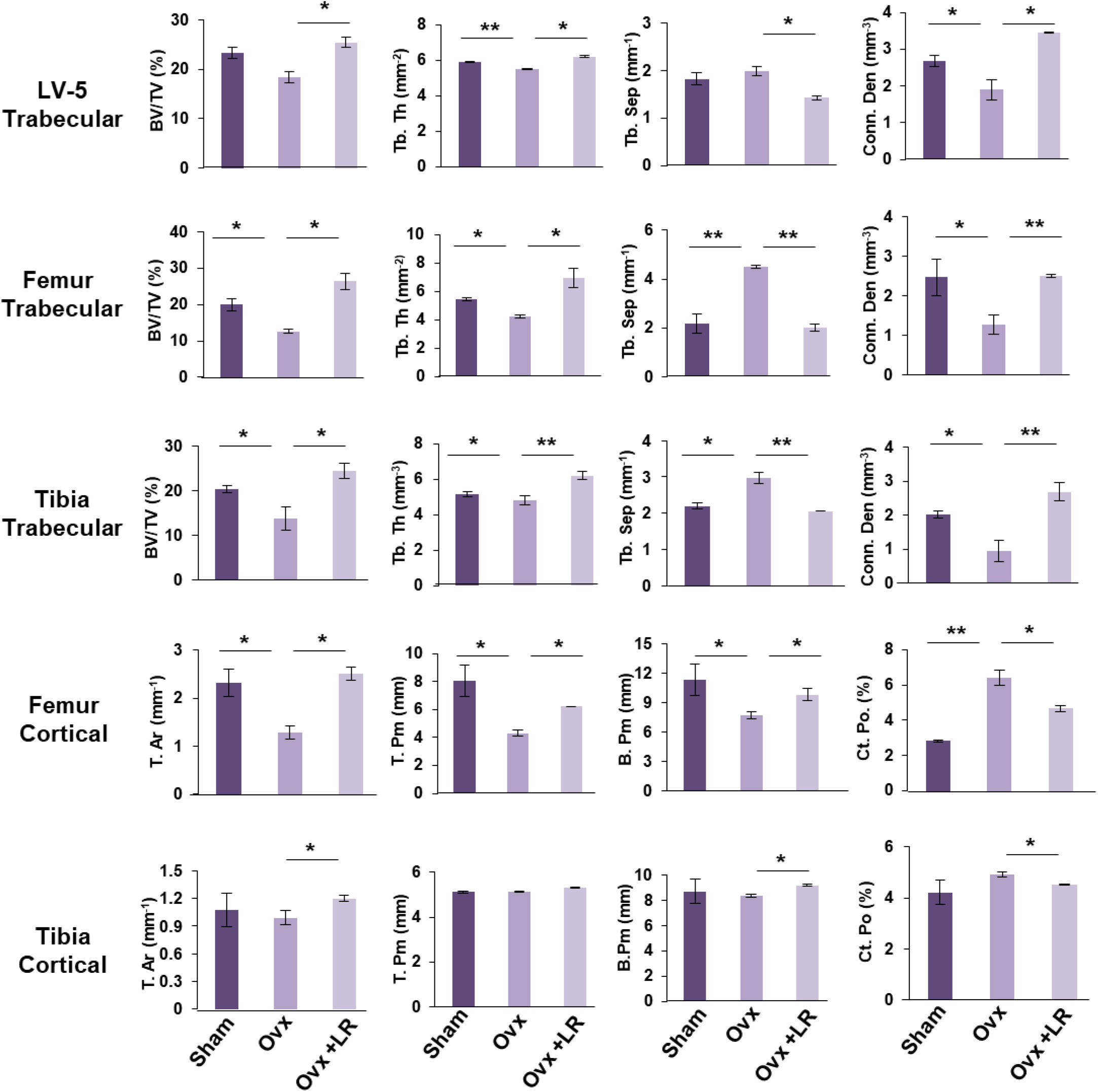
Graphical illustration of bone histo-morphometric parameters of trabecular and cortical regions. A) LV-5 Trabecular, B) Femur Trabecular, C) Tibia Trabecular, D) Femur Cortical, E) Tibia Cortical regions with different histomorphometric parameters. Bone volume/tissue volume ratio (BV/TV); Tb. Th., trabecular thickness; Conn. Den., Connective density; Tb. Sp., trabecular separation, Tt. Ar., total cross-sectional area; T. Pm., total cross-sectional perimeter; B.Pm., bone-perimeter; Ct. Po., cortical porosity. The results were evaluated by ANOVA with subsequent comparisons by Student t-test for paired or nonpaired data. Values are reported as mean ± SEM. The above graphical representations are indicative of one independent experiment and similar results were obtained in two different independent experiments with n=6. Statistical significance was considered as p≤0.05 (*p≤0.05, **p≤0.001, ***p≤0.0001) with respect to indicated mice groups.

Next, analysis of femur and tibia bones was also performed by μ-CT to analyse the effect of LR-administration on various trabecular and cortical indices in mice. Thus, μ-CT was further used to separately quantifying various bone indices parameters in femur and tibia bones. In comparison to Ovx group, 3D-micro-architecture images of trabecular and cortical regions of respective bones showed significant improvement with enhanced interconnections within the trabecular and cortical regions in Ovx+LR group (Fig. 3). Moreover, during measurement of various indices for femur trabecular region, it was found that administration of LR in Ovx mice enhanced the femur micro-architecture by augmenting the BV/TV ratio (p < 0.05), Tb.Th (p < 0.05), Conn. Den (p < 0.01) along with reducing Tb.Sp (p < 0.01) (Fig. 4). Notably, we also found comparable data in femoral cortical region with significant improvement in micro-architecture along with enhanced morphometric parameters (condition that relates to healthy bone) (Fig. 3,4). Tibial trabecular region results showed a similar trend with data clearly justifying the role of LR-administration in significantly increasing BV/TV (p < 0.05), Tb.Th (p < 0.01) and Conn.Den (p < 0.01) along with significantly decreasing Tb.Sp (p < 0.01) in Ovx mice (Fig. 4). The tibia cortical data complements our tibia trabecular results by showing higher cortical bone histo-morphometric parameters in LR-administered Ovx mice group (Fig. 4). Collectively, these results highlight the anabolic role of LR-administration on bone-health in Ovx model.

### LR elevates both mineral density and heterogeneity of bones

Mineral density of bones is a measure of mineral content packed within the segment of bones. To support our μCT data of various bones it was important to further assess the bone samples for BMD which draws clear picture about the presence of minerals and their concentration in respective bones. Bones with higher BMD are categorised as healthy and bones with lower BMD are considered as osteoporotic with higher fracture tendencies. Interestingly, we found that our BMD results complement our μ-CT results as expected, a significantly reduced BMD was found in Ovx group as compared to control group (Fig. 5). Strikingly, we observed that uptake of LR significantly enhanced and restored the BMD values of LV-5, femoral and tibial region in Ovx mice (Fig. 5a).

**Fig 5:**
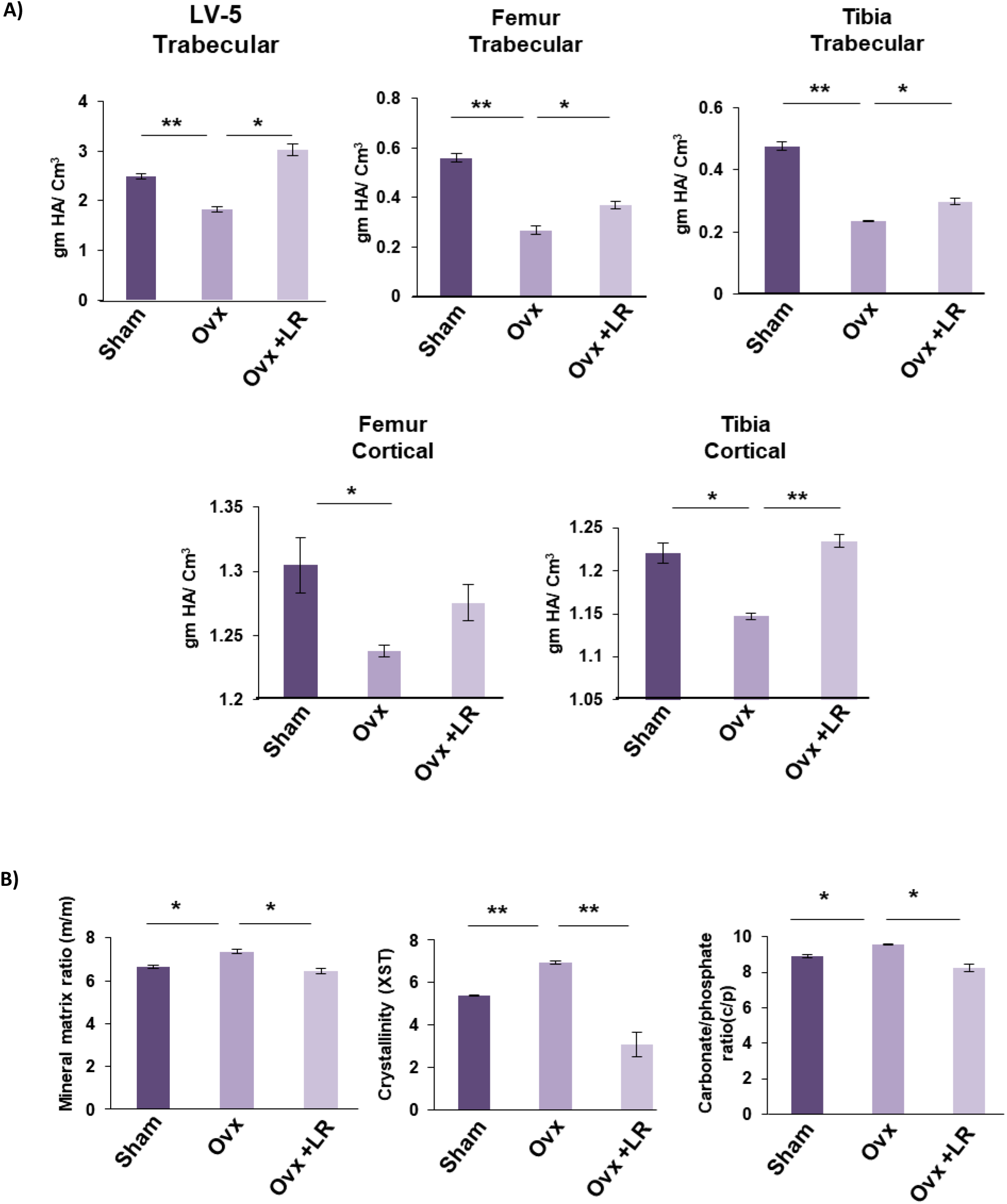
LR administration enhances both mineral density and heterogenity of bones. A) Graphical representation of BMD of LV-5, trabecular and cortical regions of femur and tibial bones of all groups. B) Graphical representation of compositional changes in bones as detected by FTIR (bone mineral/organic matrix ratio (m/m), crystallinity (XST) and carbonate to phosphate ratio (C/P). Data are reported as mean ± SEM. Similar results were obtained in two independent experiments with n=6. Statistical significance of each parameter was assessed by ANOVA followed by paired group comparison. *p < 0.05, **p < 0.01, ***p < 0.001, ****p < 0.0001 compared with indicated groups.

Composition of bones is heterogeneous and loss in bone heterogeneity has been linked with enhanced fracture risk (30). This unique feature is primarily responsible for maintaining bone physiology and functioning in a more efficient and healthy way. Fourier transform infrared spectroscopy (FTIR) happens to be at forefront among the techniques for deducing and defining structural compositional changes in organic environment including bones. Notably, it derives both qualitative and quantitative data of changes in bone compositional constituents, giving a better idea whether bones are healthy or undergoing osteoporosis (30, 31). We, therefore utilized FTIR to assess the effect of LR-administration on compositional and heterogeneity changes of bones. The analysis of bone samples revealed that Ovx mice administered with LR had significantly enhanced heterogeneity parameters viz. crystallinity (XST) (p < 0.01), carbon content (c/p) (p < 0.05) and mineral to organic matrix ratio (m/m) (p < 0.05) as compared to Ovx mice (Fig. 5b). These findings thus support and validate our hypothesis that LR-administration not only increases the BMD of bones but also preserves the natural heterogeneity of bones. Altogether, these data establish that LR may act as a suitable therapeutic probiotic in the management and treatment of osteoporotic patients.

### LR administration enhances bone health through modulation of Treg-Th17 cell balance *in vivo*

Our previous studies along with others have reported that different probiotic strains exhibit various immunomodulatory properties which are critical for inducing higher bone volume in both trabecular and cortical compartments of the skeleton (3, 14, 25). A slight disturbance in the critical balance between Treg-Th17 cells gives rise to various inflammatory conditions including osteoporosis. The balance between Tregs and Th17 immune cells is of great relevance for maintaining normal physiological functions and thus it is of great significance to decipher the role of Tregs-Th17 cells in LR regulated bone-health. Previous reports have already demonstrated that during inflammatory conditions the Treg-Th17 cell balance gets shifted towards Th17-cells, which are inflammatory in nature and thus cause enhanced bone-loss along with subsequently depleting the population of Treg-cells having osteoprotective role (14, 32, 33). Thus, we analysed the percentages of CD4^+^Foxp3^+^Tregs, CD8^+^Foxp3^+^Tregs and CD4^+^Roryt^+^Th17 immune cells in various immunological sites such as BM, spleen, PP and LN. As expected, in comparison to the sham group, the percentages of CD4^+^Foxp3^+^Tregs and CD8^+^Foxp3^+^Tregs were significantly reduced along with concurrent significant enhancement in CD4^+^Roryt^+^Th17 cell population in BM, spleen, PP and LN of Ovx group (Fig. 6). After LR-administration in Ovx mice in BM the percentages of CD4^+^Foxp3^+^Tregs were estimated to be 3-fold enhanced (p < 0.01) as compared to Ovx group treated with placebo. Interestingly, CD8^+^Foxp3^+^Tregs were also significantly enhanced (p < 0.05) along with 1.5-fold reduction in CD4^+^Roryt^+^Th17 cells in LR treated group as compared to Ovx mice (Fig. 6). In spleen, the percentages of CD4^+^Foxp3^+^Tregs were 1-fold enhanced (p < 0.01) in LR treated group as compared to placebo treated Ovx group. Strikingly, CD8^+^Foxp3^+^Tregs were also significantly enhanced (p < 0.05) along with 1.5-fold reduction in CD4^+^Roryt^+^ Th17 cells in LR treated group as compared to placebo treated Ovx group. Similarly, in LN the percentages of CD4^+^Foxp3^+^Tregs were 1-fold enhanced (p < 0.01) in LR treated group as compared to Ovx mice group. Also, CD8^+^Foxp3^+^Tregs were 3-fold enhanced (p < 0.05) in LR treated group as compared to Ovx mice with simultaneous 1.5-fold significant reduction in CD4^+^Roryt^+^Th17 cells (p < 0.05) after LR-administration in Ovx mice (Fig. 6). Further, to understand the effect of LR supplementation on lymphocytes of intestinal tissues (which are the first site of interacting with the probiotics), we examined the lymphocytes profile in PP. Strikingly, we observed 3-fold significant enhancement in CD4^+^Foxp3^+^Tregs (p < 0.001), along with 1.5-fold enhancement in CD8^+^Foxp3^+^Tregs (p < 0.01) and 1.5-fold significant reduction in CD4^+^Roryt^+^Th17 cells (p < 0.01) (Fig. 6) in Ovx mice after LR-administration. Altogether, these results suggest that LR supplementation enhances Tregs population with simultaneous reduction in Th17 cells. In accordance to previously reported studies, activated CD4^+^T cells are also known to regulate osteoclast activation by expressing RANKL on their surfaces and our results too confirmed that oral supplementation of LR to Ovx mice significantly down-regulates the expression of RANKL on CD4^+^T cells (p < 0.01) (Fig. 6). Overall, these results demonstrated that upon treatment with LR the proportion of T lymphocytes with immunosuppressive properties were significantly enhanced and the T lymphocytes exhibiting inflammatory properties were significantly reduced, thereby establishing the role of LR in modulating Treg-Th17 cell-balance in osteoporosis.

**Fig 6:**
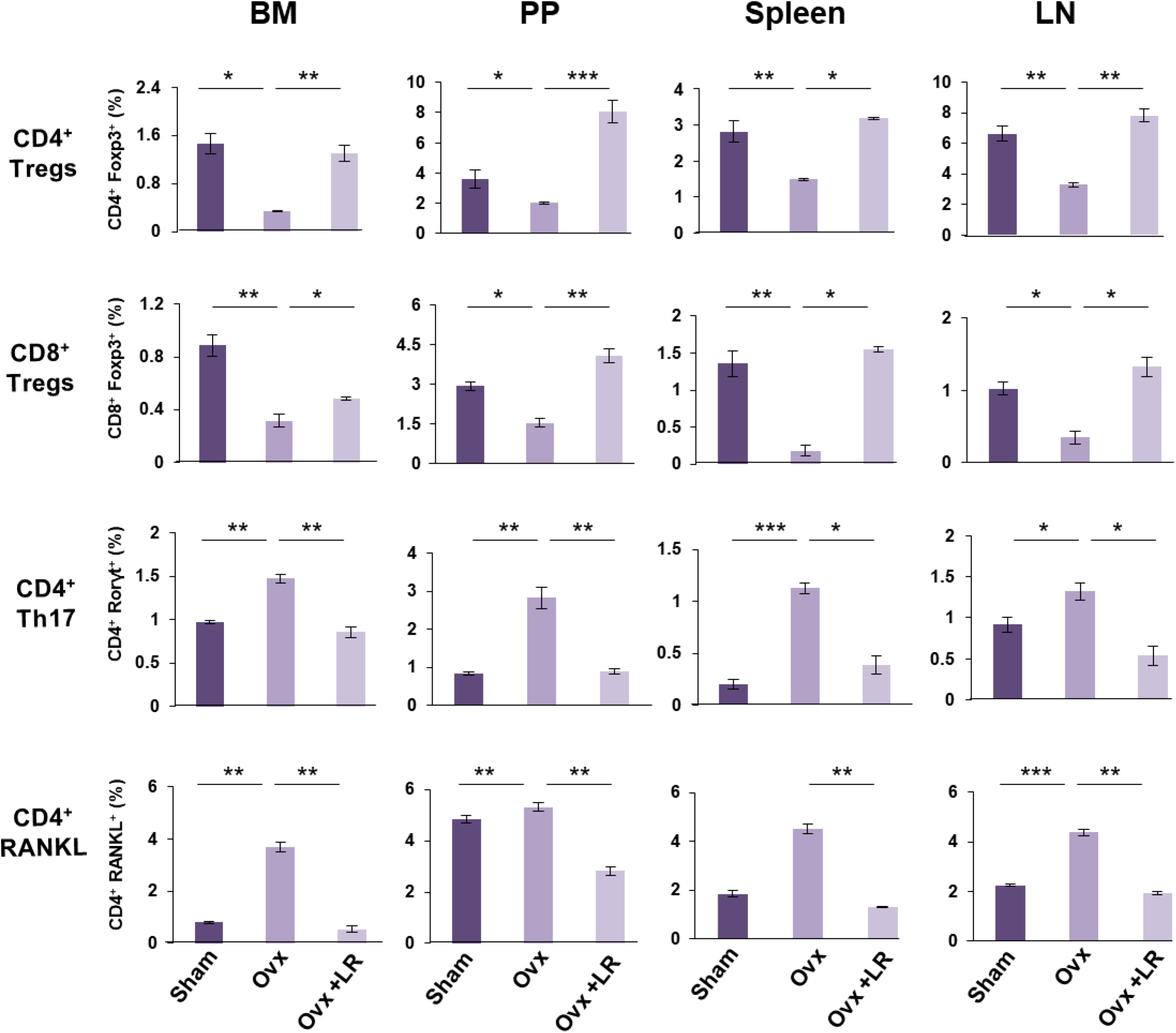
LR intake regulates Immune cells balance *in vivo*. Cells from BM, PP, spleen and LN of mice from Sham, Ovx and Ovx + LR groups were harvested and analysed by Flow Cytometry for percentage of A) CD4^+^Foxp3^+^ Tregs, B) CD4^+^Foxp3^+^ Tregs, C) CD4^+^Roryt^+^ Th17 cells D) CD4^+^ RANKL^+^ T cells. Data are reported as mean±SEM. Similar results were obtained in two independent experiments with n=6. Statistical significance of each parameter was assessed by ANOVA followed by paired group comparison. *p < 0.05, **p < 0.01, ***p < 0.001, ****p < 0.0001 compared with indicated groups.

### LR administration skews the expression of cytokines in Ovx mice

To further investigate the cellular targets of LR-administration responsible for blunting osteoclastogenesis in Ovx mice, analysis of various cytokines that plays a key role in generation and differentiation of osteoclasts in blood serum was carried out. Generally, cytokines have been categorised as either pro-inflammatory cytokines (IL-6, IL-17 and TNF-α) which enhances osteoclastogenesis and anti-inflammatory cytokines (IL-4, IL-10 and IFN-γ) which inhibit osteoclastogenesis or enhance bone formation (3, 4). A higher surge of osteoclastogenic cytokines which lead to bone-loss with a simultaneous decreased level of anti-osteoclastogenic cytokines which blunt bone-loss have been reported in post-menopausal mice model (34, 4). Thus, it was inevitable in our experiment to investigate the extent to which LR intake in Ovx mice effects expression of various cytokines. Cytokine analysis of blood serum revealed that Ovx mice administered with LR had significantly decreased levels of osteoclastogenic cytokines IL-6 (p < 0.01), IL-17 (p < 0.001) and TNF-α with significant augmentation of anti-osteoclastogenic cytokines IL-10 (p < 0.01), IL-4 (P < 0.05) and IFN-γ (0.01) (Fig. 7) as compared to Ovx-mice-group. Taken together our data strongly supports that LR-administration in Ovx-mice leads to enhanced bone-health via skewing the cytokine-balance.

**Fig 7:**
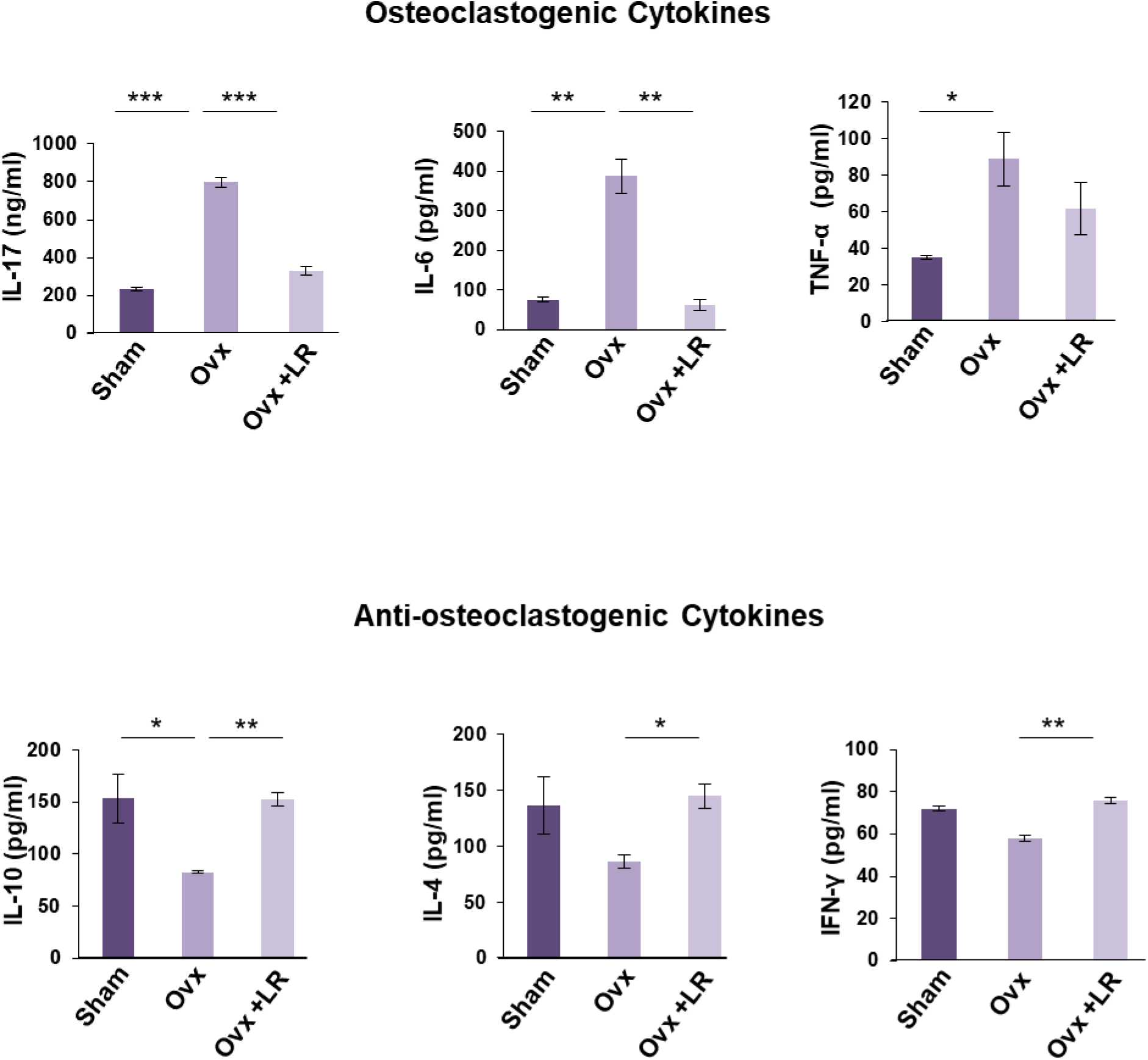
LR skews cytokines balance in ovx mice. A) Osteoclastogenic and anti-osteoclastogenic cytokines were analysed in serum samples of mice by ELISA/CBA. The results were evaluated by using ANOVA with subsequent comparisons by Student t-test for paired or non-paired data, as appropriate. Values are expressed as mean ± SEM (n=6) and similar results were obtained in two independent experiments. Statistical significance was defined as p≤0.05, *p≤0.05, **p < 0.01 ***p≤0.001 with respect to indicated mice group.

## Discussion

Disruptions in bone-remodelling process are mostly associated with elevation of osteoclast mediated bone-resorptive process that leads to exacerbation in bone-destruction (35). It has been estimated that 30% of men and 50% of women, over the age of 50 are susceptible to enhanced bone-loss and fractures. According to International-Osteoporosis-Foundation (IOF), one-third of the female and one-fifth of the male population across the globe will suffer from osteoporosis related fractures in their lifetime and it has been projected that 200 million people worldwide will suffer from osteoporosis. It has also been evaluated that the economic burden of osteoporosis by the end of 2025 will reach 25.3 billion in USA alone (36). Current therapies that are being used in adverse stages of postmenopausal osteoporosis are mainly characterized into two categories such as anti-resorptive agents and anabolic agents (37). Most of the agents that are being currently employed for treating osteoporosis are not only too costly to provide benefits to large scale population but are also associated with adverse health effects in long run (6). Thus, it is necessary to identify natural entities with no or lesser side effects that can substitute available drugs.

In recent years, growing evidences from human studies and murine models has highlighted the beneficial effects of probiotics (*Lactobacillus reuteri, Lactobacillus acidophilus, Lactobacillus casei, Bacillus clausii, Lactobacillus rhamnosus GG*) in treating various disease conditions including osteoporosis (38, 39, 40, 41, 3, 14, 42). Furthermore, the data from a recent double-blind, randomized, placebo-controlled-study have demonstrated that *Lactobacillus rhamnosus* in combination with *Lactobacillus acidophilus* and *Lactobacillus casei* reduces the symptoms and improves the quality of life of patients diagnosed with irritable-bowel-syndrome (IBS) (41). Although different studies has emphasized the significance of various probiotics including *Lactobacillus rhamnosus* in imparting beneficial effects on bone-health unfortunately the immunoregulatory role (specifically the Treg-Th17 cell-axis) of this probiotic that mediates its effects are not well understood.

Our lab has earlier shown that treatment with probiotic *Lactobacillus acidophilus* (belonging to the same genus), enhanced the femoral and tibial (both trabecular and cortical regions) bone micro-architecture, bone-mineral-content by maintaining the homeostatic balance between Tregs and Th17 cells in Ovx mice (3). A study reported by Tyagi et al. showed that administration of LR also enhanced the bone-mass in eugonadic mice (13) but the involvement of immune-cells, specifically the Treg-Th17 cell axis behind the bone-protective role of LR was not dealt in this study. Also, the effect of LR on the heterogeneity of bones and on the Tregs-Th17 homeostatic balance has not been investigated in this study. Building upon these previous evidences, in this study we have elucidated the immunomodulatory-properties by which LR maintains the bone mass in Ovx mice which mimics the postmenopausal estrogen-deficient condition in women’s.

Bone plays several crucial roles in the body such as safeguarding the internal organs, storing minerals (viz. calcium), maintaining the structure etc. thus, it is very much important to build healthy and strong bones. A condition that makes the bones brittle and weak is osteoporosis; one of the key factors involved in development of osteoporosis is alterations in hormonal levels especially in post-menopausal women, as “deficiency of estrogen” hormone makes women more susceptible to osteoporosis. Thus, to investigate the role of LR in post-menopausal osteoporosis, we developed postmenopausal-osteoporotic-mice-model (Ovx). We found that LR-administration inhibits the bone-loss in Ovx mice as confirmed by SEM and AFM data analysis of cortical bone samples which was further supported by MATLAB-data. Micro-CT analysis further reconfirmed that administration of LR significantly attenuated bone-loss by maintaining the micro-architecture of LV-5, femoral and tibial bones.

Bone strength is the best quantitative indicator of anti-fracture capability (43, 44) and is dependent on minerals present in bones. BMD of bone varies from one location of the body to other and among various locations measurement of mineral density at spine is an accurate predictor of fracture risk. BMD values narrates the risk of bone breakage or development of fracture, as higher BMD signifies lesser risk of fracture whereas lower BMD value relates to higher risk of bone breakage and fracture development. Our data outcomes evidently establish that LR has the potential of enhancing bone-health by maintaining BMDs of LV-5, femoral and tibial bones.

The nature of bone is heterogeneous due to constant remodelling process (45) this can be reflected by the distribution of mineral content, crystal composition and collagen maturity. High mineral content makes the bones more brittle whereas too low mineral content makes the bone less stiff (46). Thus, heterogeneity of bones exhibits the potential to affect the mechanical properties of whole bone (47). Crystallinity (XST) denotes the crystal-size and enhancement in XST make bones prone to fracture risk due to increased brittleness (30). Carbon to phosphate ratio (c/p) denotes the level of carbonate substitution in hydroxyapatite crystal and it has been reported that c/p ratio increases in osteoporotic bones (48, 30, 14). In consistent to this our study too demonstrated that as compared to the Ovx group the m/m ratio, crystallinity and c/p ratio were significantly reduce in Ovx-group treated with LR. Thus, corroborating the fact that LR-administration regulate mechanical property of bones by maintaining the natural heterogeneous composition of bones.

Both bone cells and immune cells shares a common niche (i.e. bone-marrow) during their development, an active field of research termed as Osteoimmunology. Among other immune cells, Tregs and Th17 lymphocytes are the key players involved in regulating the bone-remodelling process and exert effect on both the bone forming osteoblasts and bone resorbing osteoclasts. Cytokines derived from these immune cells such as IL-10 and IL-17A have established roles in regulating both osteoclastogenesis as well as osteoblastogenesis (49, 50, 51, 52, 4). In this context, we too studied the effect of probiotic LR on Tregs-Th17 cells modulation in BM (prime-site-of-osteoclastogenesis), spleen and LN and our data clearly suggests that LR exerts systemic bone effects via enhancing Tregs population along with simultaneous reduction in Th17 cells. Since the GUT harbours more than 70% of the host immune system (53). Also, the intestinal tissue provides residence to various immune cells found to be in consistent interaction with the intestinal microbiota. Probiotics such as *Lactobacillus reuteri* via its immunomodulatory properties mediates its beneficial effects on bone-health by modulating immune cells in intestinal tissue. With respect to this, we also investigated the role of Tregs-Th17 population residing in gut lymphoid organ i.e. PP. We observed that LR-administration modulated the Treg-Th17 cell axis in PP towards the Tregs lineage and thus improved bone-health in Ovx mice. These results were further supported by the serum cytokine analysis data where we observed enhancement of anti-osteoclastogenic cytokines such as IL-4, IL-10 and IFN-γ that exhibits potential to reduce osteoclastogenesis (9) and down regulation of inflammatory cytokine levels such as IL-6, IL-17 and TNF-α exhibiting property to enhance osteoclastogenesis. LR-administration also hampers the RANKL mediated osteoclastogenesis by suppressing the expression of RANKL on CD4^+^T cells at distinct sites. Thus, these data suggest that LR enhances bone-health via maintaining the homeostatic balance between Tregs-Th17 cells (Fig. 8). Altogether our data further support the existing facts that probiotics can impart beneficial-effects on health via exhibiting immunomodulatory properties (3, 14, 25).

**Fig 8:**
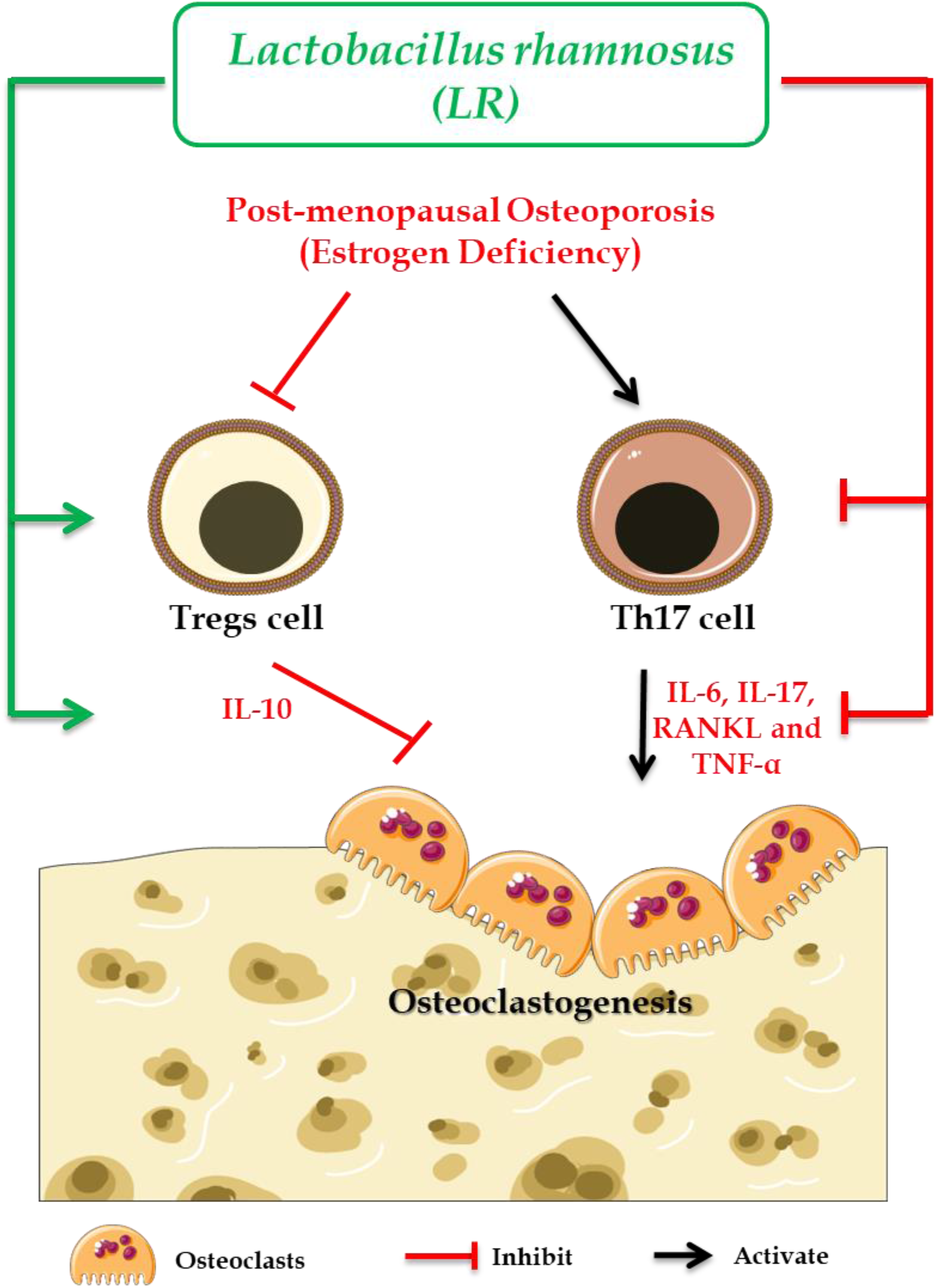
Summary of our results. LR administration attenuates bone loss via modulating the Tregs-Th17 balance and its secretary cytokines in Ovx mice. (Image illustrated using Medical Art https://smart.servier.com/).

Our present study thus for the first time demonstrated the beneficial effects of LR on skeletal health by maintaining the BMD, preserving the micro-architecture and heterogeneity of bones in bilaterally-induced-ovariectomized-mice-model via modulating the homeostasis between Tregs and Th17 cell axis in the host. The present study thus highlights the potential of probiotic LR to be used as a novel osteo-protective-agent in the treatment and management of bone-related-diseases including osteoporosis.

## Materials and Methods

### Animals

All *in vivo* experiments were performed in 8-10-wks-old-female BALB/c SPF mice. Mice were maintained under specific pathogen free conditions at the animal facility of All India Institute of Medical Sciences (AIIMS) New Delhi, fed sterilized food and autoclaved-drinking-water *ad-libitum*. Mice were exposed to bilateral-ovariectomy (Ovx) and sham surgery after anesthetizing them with ketamine (100-150mg/kg) and xylazine (5-16mg/kg) intraperitoneally. Later, operated mice were divided into three groups with 6 mice in each group; group A: sham operated; group B: Ovx + placebo; group C: Ovx + *Lactobacillus rhamnosus* (LR). *Lactobacillus rhamnosus* UBLR-58 (MTCC 5402) was procured from Unique-Biotech-Ltd., Hyderabad, India.

After one-week post-surgery, LR was administered orally as suspension of 400ul (10^9^ cfu/ml) daily dissolved in drinking water to Ovx+LR group for a period of 6wks and the same volume of placebo was also given to Ovx+placebo group. At the end of experiment (6wks), animals were euthanized by carbon-dioxide-asphyxiation and blood, bones and lymphoid tissues were harvested for further analysis (Fig. 1A). The body weight of animals was recorded at regular intervals during the experimental period. All the procedures were performed in accordance with the principles, recommendation and after due approval of the protocols submitted to Institutional Animal Ethics Committee of All India Institute of Medical Sciences (AIIMS) (71/IAEC-1/2018).

### Reagents and Antibodies

Below mentioned antibodies/kits were acquired from eBiosciences (USA): APC-Anti-Mouse/Rat Foxp3-(FJK-16s)-(17-5773), PE-Anti-Human/Mouse-Rorγt-(AFKJS-9)-(12-6988), Foxp3/Transcription-factor staining-buffer (0-5523-00), RBC-lysis-buffer (00-4300-54), Following ELISA kits were brought from R&D: Mouse-IL-10-(M1000B) and Mouse-IL-17-(M1700)-Quantikine ELISA-kits. Following ELISA kits were brought from BD: Mouse-IL-6 (OptEIA™-555240) and Mouse-TNF-α-(OptEIA™-560478). (PerCp-Cy5.5-Anti-Mouse-CD4-(RM4-5)-(550954) was obtained from BD-Biosciences. PE/Cy7-Anti-Mouse-CD8-(53-6.7)-(100721), PE-Anti-Mouse-CD254-(TRANCE-RANKL)-(IK22/5)-(510005) were acquired from Biolegend (USA).

### Scanning Electron Microscopy (SEM)

SEM for femur cortical region of bones was done as described previously (3, 14, 29). Briefly, bone samples were stored in 1%-Triton-X-100 for 2-3 days and later bone-samples were transferred to 1XPBS buffer till the final analysis was performed. After preparation of bone slices, samples were dried under incandescent lamp and sputter coating performed. Subsequently bones were scanned in Leo 435-VP-microscope equipped with digital-imaging with 35 mm photography system. SEM-images were digitally-photographed at 100X-magnification to capture the best-cortical-regions. The SEM-images were further analysed by MATLAB-software (Mathworks, Natick, MA, USA).

### Atomic Force Microscopy (AFM)

After drying femur bone samples completely in sterile environment with 60W lamps for 6 hours followed by high vacuum drying. Samples were prepared as per requirement for the machine and analysed by Atomic Force Microscope (INNOVA-ICON, Bruker) that works under the Acoustic-AC-mode. This was assisted by cantilever (NSC 12(c) MikroMasch, Silicon-Nitride-Tip) and NanoDrive™ version-8-software. It was set at constant force of 0.6N/m with a resonant frequency at 94-136 kHz. Images were recorded at a scan-speed of 1.5-2.2 lines/s in air at room-temperature. Images were later-processed and analysed by using nanoscope-analysis-software. Further, the 3D AFM images were also analysed by MATLAB-software (Mathworks, Natick, MA, USA).

### Micro-Computed-Tomography (μ-CT) measurements

μ-CT scanning and analysis was performed as described previously (3, 14, 29) using SkyScan 1076 scanner (Aartselaar, Belgium). Briefly, after positioning all samples at right orientation, scanning was done at 50 kV, 201mA using 0.5 mm aluminium filter and exposure was set to 590 ms. NRecon-software was used for carrying out reconstruction process. For trabecular region analysis, ROI was drawn at a total of 100 slices in secondary spongiosa at 1.5mm from distal border of growth plates excluding the parts of cortical bone and primary spongiosa. The CTAn-software was used for measuring and calculating the micro architectural parameters in bone samples. Various 3D-histomorphometric-parameters were obtained such as: BV/TV (Bone volume/Tissue volume), Tb.Th (Trabecular-thickness), Tb.Sp. (Trabecular-separation) and Conn.Den (Connective-density) etc. The volume of interest of u-CT scans made for trabecular and cortical regions were used to determine the bone-mineral-density (BMD) of LV5, femur and tibia. BMD was measured by using hydroxyapatite phantom-rods of 4 mm diameter with known BMD (0.25 g/cm3 and 0.75 g/cm3) as calibrator (3, 14, 29).

### Fourier Transform Infrared (FTIR) Spectroscopy

Femur cortical bone samples were stored in 1% Triton X-100 for 24 hours before being dried with 60W lamps for 6 hours followed by high vacuum drying. Dried bone samples were crushed in pestle-mortar, thereafter bone-samples were mixed with potassium-bromide (KBr) at (1:100) ratio for FTIR-analysis. Further, acquisition was performed by using 8400S-FTIR- (SHIMADZU), with a resolution 4cm^−1^; scan speed 2.5 kHz and 128 scans. The samples were clearly positioned with a prism made of highly refractive material. Savitzky-Golay-algorithm was used to nullify background noise for obtaining smooth spectra of all analysed-samples (3, 14, 29).

### Flow Cytometry

Experiment was carried out as per the standard-protocol mentioned previously (3). Briefly, after preparing single cell suspension of BM, PP, spleen and LN, RBC-lysis was performed by RBC-lysis buffer and targeted cell population was stained with specific antibodies labelled with particular fluorochromes. For surface marker staining, cells were first incubated with cocktail of cell surface antibodies: Anti-CD4 and Anti-CD8 antibodies for 30’ in dark on ice. After washing with 1X wash buffer, cells were fixed with 1X fixation-permeabilization-buffer for 30’ on ice in dark. Further for intracellular staining, cells were washed with 1X permeabilization-buffer and stained with the following cocktail of antibodies: Anti-Rorγt, Anti-Foxp3 and Anti-RANKL for 45 minutes on ice. Washing was done by using permeabilization-buffer and cells were acquired on BD-LSR-II flow-cytometer. Flojo-10 (TreeStar, USA) software was used to analyse the samples and gating strategy was done as per previously defined-protocols (14).

### Enzyme Linked Immunosorbent Assay (ELISA) and Cytometric Bead Array (CBA)

ELISA was performed for quantitative assessment of cytokines viz. IL-4, IL-6, IL-10, IL-17 and TNF-α in blood serum using commercially available kits. For estimating IFN-γ cytokine in serum, CBA was performed as per the manufacturer’s instructions (BD-Biosciences). Fluorescent signals were read on flow analyser and data analysed by BD-FCAP-Array software (BD-Biosciences).

### Statistical Analysis

Statistical differences between groups were assessed by using analysis-of-variance (ANOVA) with subsequent comparisons via student t-test paired or unpaired as appropriate. All the data values are expressed as Mean±SEM (n=6). Statistical significance was determined as P ≤ 0.05 (*p < 0.05, **p < 0.01, ***p < 0.001, ****p < 0.0001) with respect to indicated group.

## Acknowledgment

This work was financially supported by projects: DST-SERB (EMR/2016/007158), Govt. of India and intramural project from All India Institute of Medical Sciences (AIIMS, A-596), New Delhi-India sanctioned to RKS. HYD, LS, AP, ZA, SK, AB and RKS acknowledge the Department of Biotechnology, AIIMS, New Delhi-India for providing infrastructural facilities. HYD thank ICMR research fellowship, LS and ZA thank UGC for research fellowship, AP thanks thank DBT for PG fellowships and AB thank DST SERB project for research fellowship.

## Author contributions

RKS contributed in conceptualization and investigation of study. HYD, AP, LS, ZA, SK and AB contributed in methodology. RKS, HYD, LS and AP contributed in formal analysis of data. PS analysed SEM and AFM data for MATLAB analysis. PKM carried out cytokine analysis. LS, HYD and RKS contributed in writing and editing of manuscript. BV provided valuable inputs in the study design. All authors reviewed the manuscript.

## Compliance with ethical standards

All applicable institutional and/or national guidelines for the care and use of animals were followed.

## Abbreviations

IFN: interferon
IL: interleukin
TNF: tumor necrosis factor
RANKL: receptor activator of nuclear factor kappa β ligand
Ovx: ovariectomized
BMD: bone mineral density
BM: bone marrow
PP: peyer’s patches
LN: lymph nodes
RA: rheumatoid arthritis
2D: two dimensions
3D: three dimensions.

